# Productive infection of field strains of avian coronavirus infectious bronchitis virus in chicken peripheral blood-derived monocytes

**DOI:** 10.1101/041558

**Authors:** Yueting Zhang, Gary R. Whittaker

## Abstract

The avian coronavirus infectious bronchitis virus (IBV) typically infects the respiratory tract of chickens, but can also spread to other organs. However, the mechanisms of virus dissemination are presently unclear. We show that peripheral blood-derived monocytes/macrophages from chickens (chPBMC) are productively infected by clinical strains of IBV, accompanied by induction of apoptosis. Our data suggest that chPBMCs play a role in the dissemination of IBV, and may be important for viral pathogenesis.

## Introduction

Avian infectious bronchitis virus (IBV) is the prototype member of the Gammacoronavirus genus (1, 6). Despite extensive vaccination, IBV poses a continued economic threat to the poultry industry. The primary site of IBV infection is the epithelium of the respiratory tract; however IBV can spread to other sites including the gastrointestinal tract, kidney reproductive organs and Harderian glands (2, 9). Viremia has been suggested as a way for internal spread, as evidenced by detection of virus in the blood of infected chickens (3, 5). However, the mechanism remains unknown. Other coronaviruses, such as severe acute respiratory syndrome-coronavirus (4) and feline coronavirus (10), exhibit well recognized systemic infection, with monocytes, macrophages and dendritic cells originating from a common myeloid progenitor disseminating the virus and acting as an important immunopathogenic component in disease (8).

In the case of IBV, previous studies have shown that chicken bone marrow-derived macrophages were resistant to infection *in vitro* (11). However, macrophages from different anatomical sites can elicit distinct functions or responses to foreign pathogens (7, 8). To explain the known presence of IBV in the blood of infected chickens (3, 5), we reasoned that peripheral blood-derived monocytes/macrophages from chickens (chPBMCs) could be susceptible to IBV infection.

## Methods

IBV strains used in this study were M41, Cal99, Conn46, Iowa97 (all field strains) and Beaudette (a laboratory-adapted version of M41). Virus stocks were produced in embryonated chicken eggs and titered by EID50 or TCID50 assay. Chicken peripheral blood monocytes were prepared as described (20). Briefly blood was obtained by cardiac puncture from 6-8 week-old specific-pathogen-free White Leghorn chickens. Monocytes were isolated with a Ficoll gradient (GE Healthcare) and were cultured in RPMI 1640 supplemented with 10% heat-inactivated chicken serum and penicillin/streptomycin. 24h after cell seeding non-adherent were washed away. Monocytes for each independent experiment were obtained from an individual blood donor. All work with chickens was carried out according to the Cornell University Animal Care and Use program and complied with the Public Health Service Policy on Humane Care and Use of Laboratory Animals

For immunofluorescence assay, cells were fixed with methanol for detection of IBV viral antigens; otherwise cells were fixed with paraformaldehyde. Anti-S1 monoclonal antibody 15:88 was used for IBV-M41 and Beaudette, and anti-M monoclonal antibody 9:19 was used for IBV-Cal99, IBV-Conn46, and IBV-Iowa97. Monoclonal antibody KUL01 was purchased from Abcam. Secondary antibody was AlexaFluor goat anti-mouse 488 and nuclei were stained with Hoechst 33258 (Molecular Probes). Cells were quantified by scoring the % cells positive for viral antigen with >200 cells quantified from three independent experiments.

For RT-PCR for viral detection, 106 PBMCs were infected with 7x102 TCID50 IBV-M41. At indicated time points, culture supernatant was collected and filtered. Viral RNA was extracted from the supernatant using a Qiagen Viral RNA extraction kit. The RT-PCR reaction used a SuperScript One-Step RT-PCR kit (Invitrogen). IBV N gene primers were previously described by (17).

For apoptosis assays, chPBMCs were infected with 2x10^3^ TCID_50_IBV-M41 and at 6h post infection, we used a Green FLICA caspase 3/7 assay kit (ImmunoChemistry Technologies). For the DNA laddering assay, total DNA was extracted from chPBMCs using a Qiagen DNeasy Blood & Tissue kit, and laddering was visualized on an agarose gel. To inactivate IBV-M41, viruses were irradiated for 30min under a UV-lamp (wavelength 254nm).

## Results and Discussion

### Peripheral blood-derived monocytes from chickens are infected by IBV

To investigate a role for myeloid cells in IBV dissemination and viral pathogenesis, we isolated peripheral blood-derived monocytes from chickens (chPBMCs) and confirmed that they were positive for the monocyte/macrophage marker KUL01 (Fig.1A). Due to the lack of available reagents for avian species, we were unable to distinguish whether chPBMCs were matured or differentiated into macrophages or dendritic cells, or to determine the composition of unique cell populations. We found that chPBMCs are highly susceptible to the IBV field strains M41, Cal99, Conn46, and Iowa97 (Fig.1B). However, the laboratory-adapted strain Beaudette, which is not pathogenic in chickens, did not infect chPBMCs. Infections were also repeated with macrophages isolated from bone marrow of chickens, with similar results (not shown).

**Figure 1.**
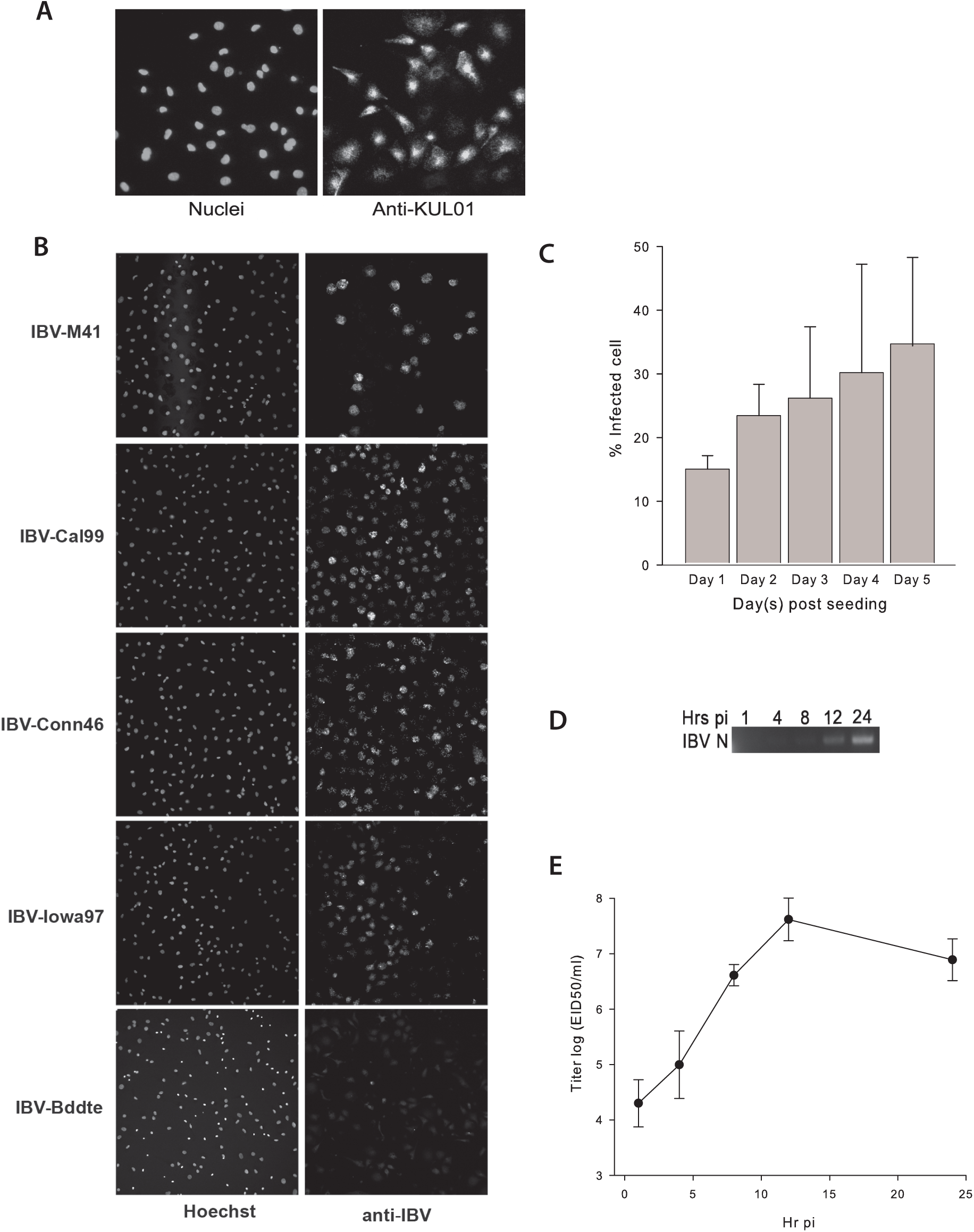
Chicken PBMCs can be productively infected by field strains of IBV. (A) chPBMCs (3-day-old culture) were labeled with mouse anti-KUL01 avian macrophage/monocyte marker antibody followed by incubation with secondary antibody goat anti-mouse. Cell nuclei were stained with Hoechst 33258. (B) chPBMCs were infected with IBV-M41, IBV-Cal99, IBV-Conn46, IBV-Iowa97 or IBV-Bddte for 1 h at 37°C. Cells were washed with PBS and further incubated for 9 h for IBV-M41 and IBV-Bddte and for 23 h for other IBVs. Viral spike protein antigen was detected by immunofluorescence microscopy. Cell nuclei were stained with Hoechst 33258. (C) Infectivity of IBV-M41 in day 0 to 5 cultures of chPBMCs was determined by immunofluorescence microscopy. Cells were quantified by scoring the percentage of cells positive for viral antigen S or M. >200 cells were quantified from each of three independent experiments. (D) Filtered supernatants were harvested from IBV-M41 infected chPBMC at indicated time points. RT-PCR against IBV N gene was performed and amplified DNA products were visualized on an agarose gel. (E) The infectious titer of the supernatants containing IBV particles was determined by EID50 assay.

To examine whether the stage of maturation of the chPBMCs had an effect on IBV infection, we tested chPBMCs at different days post-seeding (Fig.1C). The percentage of infected cells increased with time, reaching a maximum at day 5. Beyond day 3-4, we observed a large proportion of giant multinucleated cells, as observed previously (12). In all further experiments, we utilized day 3-4 cultures due to difficulty in imaging and quantifying the giant cells.

We next determined whether IBV infection of chPBMCs produces progeny viral particles. RT-PCR was performed to detect released particles. Starting at 8h, we detected viral RNA in the supernatant, reaching a maximum at 24h postinfection. To determine whether these progeny virions were infectious, we performed an EID50 assay. The peak viral titer was 10^8^ EID_50_/mL at 12h post-infection. At later times, the titer decreased slightly even though RT-PCR indicated more virus particles were present (Fig.1D). The reduced infectivity may be due to aggregation of viral particles between 12 and 24 h post infection.

### IBV induces rapid apoptosis in chPMBCs

One feature of chPBMC infection that we wished to examine was whether IBV can induce apoptosis. To do this, we used a FLICA assay that employs an inhibitor irreversibly binding to activated caspase3/7. At 6h post-infection, there was an increase in the fluorescence signal in IBVM41-infected cells, compared to mock-infected cells. Addition of UV-inactivated IBV-M41 also resulted in an increase in fluorescence signal, indicating that replication-defective IBV is also able to trigger apoptosis (Fig.2A). Cellular DNA from mock infected or infected chPBMCs was also collected after 24h post-infection and analyzed by gel electrophoresis. DNA from infected cells was fragmented and displayed an apoptosis-characteristic laddering while no obvious laddering was observed for mock-infected cells (Fig.2B).

**Figure 2.**
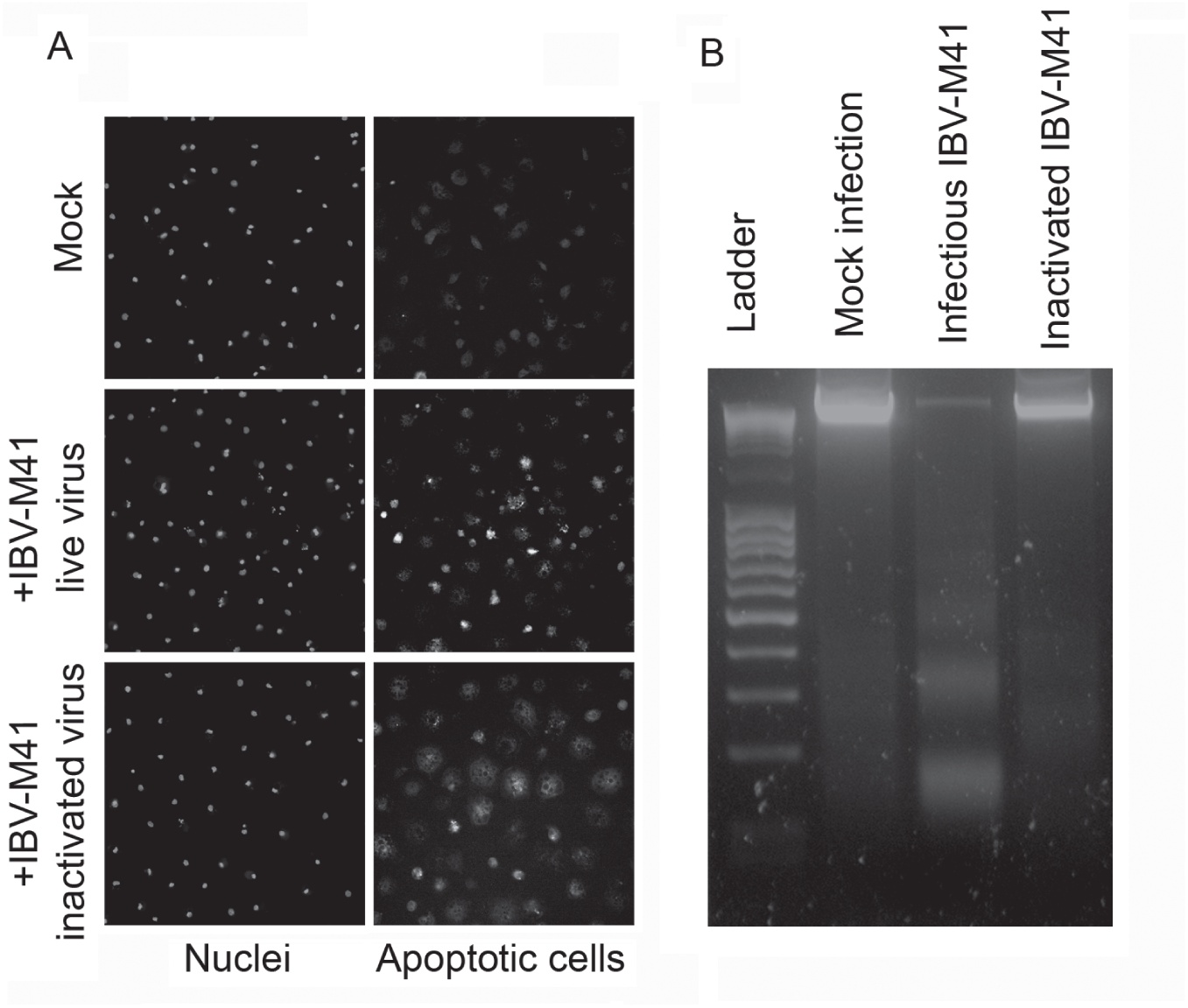
IBV-M41 induces rapid apoptosis in chPBMCs. (A) Mock infected, live IBV-M41, or inactivated IBV-M41 infected chPBMCs were assayed at 6 h post infection. Apoptosis was revealed by a fluorescence FLICA caspase 3/7 detection assay. (B) Mock infected, IBV-M41(5x10^2^TCID_5_0) infected, or UV-inactivated IBV-M41(5x10^2^TCID_5_0) infected chPBMCs were harvested and total cellular DNA extracted and analyzed on an agarose gel.

We show here that field strains of IBV can productively infect monocytes/macrophages and induce apoptosis, yet the laboratory-adapted strain Beaudette does not infect. Our data suggest that IBV infection of blood-derived myeloid cells might be an important component in the pathogenesis of IBV, and be critical for viral dissemination.

## Acknowledgments

We thank Nadia Chapman and Douglas Haner for their excellent technical support, Jean Millet and Long Ping Victor Tse for critical reading of this manuscript and all members of Whittaker laboratory for helpful discussions. YZ was supported by grant T32AI007618 (Training in Molecular Virology and Pathogenesis) from the National Institutes of Health. This work was funded by a research grant from the Cornell College of Veterinary Medicine

